# The role of auxiliary parameters in evaluating voxel-wise encoding models for 3T and 7T BOLD fMRI data

**DOI:** 10.1101/2020.04.07.029397

**Authors:** Moritz Boos, J. Swaroop Guntupalli, Jochem W. Rieger, Michael Hanke

## Abstract

In neuroimaging, voxel-wise encoding models are a popular tool to predict brain activity elicited by a stimulus. To evaluate the accuracy of these predictions across multiple voxels, one can choose between multiple quality metrics. However, each quality metric requires specifying auxiliary parameters such as the number and selection criteria of voxels, whose influence on model validation is unknown. In this study, we systematically vary these parameters and observe their effects on three common quality metrics of voxel-wise encoding models in two open datasets of 3- and 7-Tesla BOLD fMRI activity elicited by musical stimuli. We show that such auxiliary parameters not only exert substantial influence on model validation, but also differ in how they affect each quality metric. Finally, we give several recommendations for validating voxel-wise encoding models that may limit variability due to different numbers of voxels, voxel selection criteria, and magnetic field strengths.

## Introduction

In functional magnetic resonance imaging (fMRI) research, voxel-wise encoding models are an increasingly popular tool to characterize the relationship between a real world stimulus and BOLD activity patterns [Thirion et al., 2006, Kay et al., 2008, Naselaris et al., 2011, Huth et al., 2012, Holdgraf et al., 2016]. However, researchers face many degrees of freedom in constructing voxel-wise encoding models, such as how to represent the stimulus or how to estimate an accurate model. To evaluate the fidelity of these choices and the resulting voxel-wise encoding models, multiple quality metrics exist that assess how well these models approximate spatial patterns of brain activity across multiple voxels. Mitchell et al. [2008] first introduced binary retrieval accuracy - a measure that assesses whether an encoding model’s predicted spatial activation patterns can, on average, identify the correct stimulus against a decoy - to evaluate the predicted fMRI images for the meaning of nouns. Kay et al. [2008] used stimulus identification accuracy, and Naselaris et al. [2009] used stimulus reconstruction quality as a metric for encoding performance. In auditory neuroscience, only binary retrieval accuracy [Casey et al., 2012, Hoefle et al., 2018] and stimulus identification [Santoro et al., 2014, Allen et al., 2018] have been used. However, while providing novel insights into the cortical processing of sensory stimuli, it is unknown how additional parameters of the data analysis that are unrelated to the encoding models themselves affect these quality metrics. Hence, it remains difficult to compare the quality of voxel-wise encoding models when they differ in such auxiliary parameters. In this study, we aim to tackle this problem by documenting the effect of two common auxiliary parameters in the validation of voxel-wise encoding models - the number of voxels used and how they were selected - on different quality metrics for low and high field strength (3 and 7-Tesla fMRI). To this end, we utilize a 3T fMRI study on the perception of musical genres [Casey et al., 2012] which has recently been replicated in 7T [Hanke et al., 2015]. Using these datasets, we apply three approaches to encoding model validation, - binary retrieval accuracy [Mitchell et al., 2008], stimulus identification [Kay et al., 2008, Santoro et al., 2014] and decoding accuracy of the stimulus [Naselaris et al., 2009] -, and compare them in a 3T and 7T dataset with identical stimuli and comparable design, for different choices in data analysis parameters.

## Methods

### Stimuli

Stimuli were five natural, stereo, high-quality music stimuli (6 s duration; 44.1 kHz sampling rate) for each of five different musical genres: 1) Ambient, 2) Roots Country 3) Heavy Metal, 4) 50s Rock’n’Roll, and 5) Symphonic. Previously, all 25 stimuli have been made publicly available [Hanke et al., 2015].

### fMRI data

The analyses presented here were performed on two independently recorded, and previously published datasets [Casey et al., 2012, Hanke et al., 2015]. While these datasets have been acquired using identical stimuli, with the same number of acquisition runs and number of stimulation trials, they nevertheless differ in their precise stimulation timing, stimulation setup and equipment, as well as other acquisition details. A brief description of both datasets is provided below. For more information the reader is referred to the respective publications.

#### 3 Tesla

Participants were scanned in a Philips Intera Achieva scanner with 32 channel SENSE head coil at the Center for Cognitive Neuroscience at Dartmouth College. Functional scans were acquired with an echo planar imaging sequence (2 s TR; 35 ms TR, 90° flip angle) with 3 mm isotropic voxels. Each participant participated in eight functional runs. Stimuli were presented in an event-related design with a variable trial duration. Each run consisted of a total of 29 trials corresponding to 25 music clips and 4 catch trials presented randomly during each run. Each trial started with a 6 s music clip followed by 4-8 s of fixation. For catch trials, a question appeared after the audio presentation asking whether an acoustic feature is present in the music clip such as vocals, guitar, etc. Subjects responded “Yes” or “No” with a button box. Catch trials were supposed to help keep the participants’ attention to the music and were discarded from the analyses. Each run had 4 s of fixation at the beginning and 10 s of fixation at the end. For further details, see Casey et al. [2012]

#### 7 Tesla

The procedures for the 7 Tesla acquisition were highly similar and only criticial differences are reported here. Echo-planar BOLD images (gradient-echo, 2 s repetition time (TR), 22 ms echo time, 0.78 ms echo spacing, GRAPPA acceleration factor 3) were acquired using a whole-body 7 Tesla Siemens MAGNETOM magnetic resonance scanner equipped with a 32 channel brain receive coil. 36 axial slices (thickness 1.4 mm, 1.4 *×* 1.4 mm in-plane resolution) with a 10% inter-slice gap were recorded in ascending order. Slices were oriented to include the ventral portions of frontal and occipital cortex while minimizing intersection with the eyeballs. The field-of-view was centred on the approximate location of Heschl’s gyrus along the rostral-caudal axis.

Instead of dedicated catch trials, similar catch questions as for the 3T acquisition were presented 4 s after the end of the stimulus in trials with an 8 s inter-stimulus delay. Consequently, each run consisted of 25 trials, and no trials were discarded from the analysis. There was no additional fixation at the start of a run. For further details, see Hanke et al. [2015], and Hanke et al. [2014] for details on MRI acquisition methods.

### Preprocessing

Approximate temporal lobe masks for each participant were extracted from Montreal Neurological Institute coordinate space using FSL [Smith et al., 2004, Jenkinson et al., 2012], and projected into the subject-specific coordinate system. Each voxel inside the temporal lobe mask was run-wise *Z*-scored and linearly de-trended using PyMVPA [Hanke et al., 2009]. After preprocessing, 3T fMRI data consisted of 16999 voxels and 7T fMRI data of 115439 voxels.

### Encoding model

To build an encoding model with high predictive power, one needs to find an appropriate feature representation of the music stimuli [Holdgraf et al., 2017]. Casey et al. [2012] compared different feature representations of the same stimuli in the 3T dataset. They found features corresponding to the timbre of the stimulus offer the best discriminative power. We chose a similar feature set, related to timbre, that was made available by Hanke et al. [2015], the low-quefrency mel-frequency spectrum (LQ-MFS) of the stimulus. This stimulus representation consists of 48 LQ-MFS coefficients for each 100 ms segment of a stimulus, thus each stimulus overall comprises 2880 LQ-MFS coefficients.

For each fMRI sample *y*_*vt*_ (where *t* = 1, 2,, …, *T* denotes the time-points, *T* the number of samples in each run, and *v* = 1, 2, …, *V* denotes the voxels) the LQ-MFS features 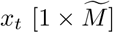 (where 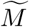 is the number of LQ-MFS coefficients) of the corresponding two second part of the stimulus - immediately prior to the acquision time point - were computed. In case there was no stimulus presented at time-point *t*, a zero vector 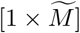 was used.

As the BOLD response is delayed, the most recent feature vector was removed for each fMRI sample (this corresponds to an assumed 2 s stimulus-response delay), and the new feature vector at time-point *t* was created by concatenating the prior feature vectors *x*_*t−*1_,*x*_*t−*2_ and *x*_*t−*3_ (see Figure 1). From now on, we denote this stacked feature vector as *x*_*t*_. Feature vectors (and the corresponding fMRI sample) were removed from the analysis, if two-thirds or more of the concatenated feature vectors were zero-vectors.

**Figure 1:**
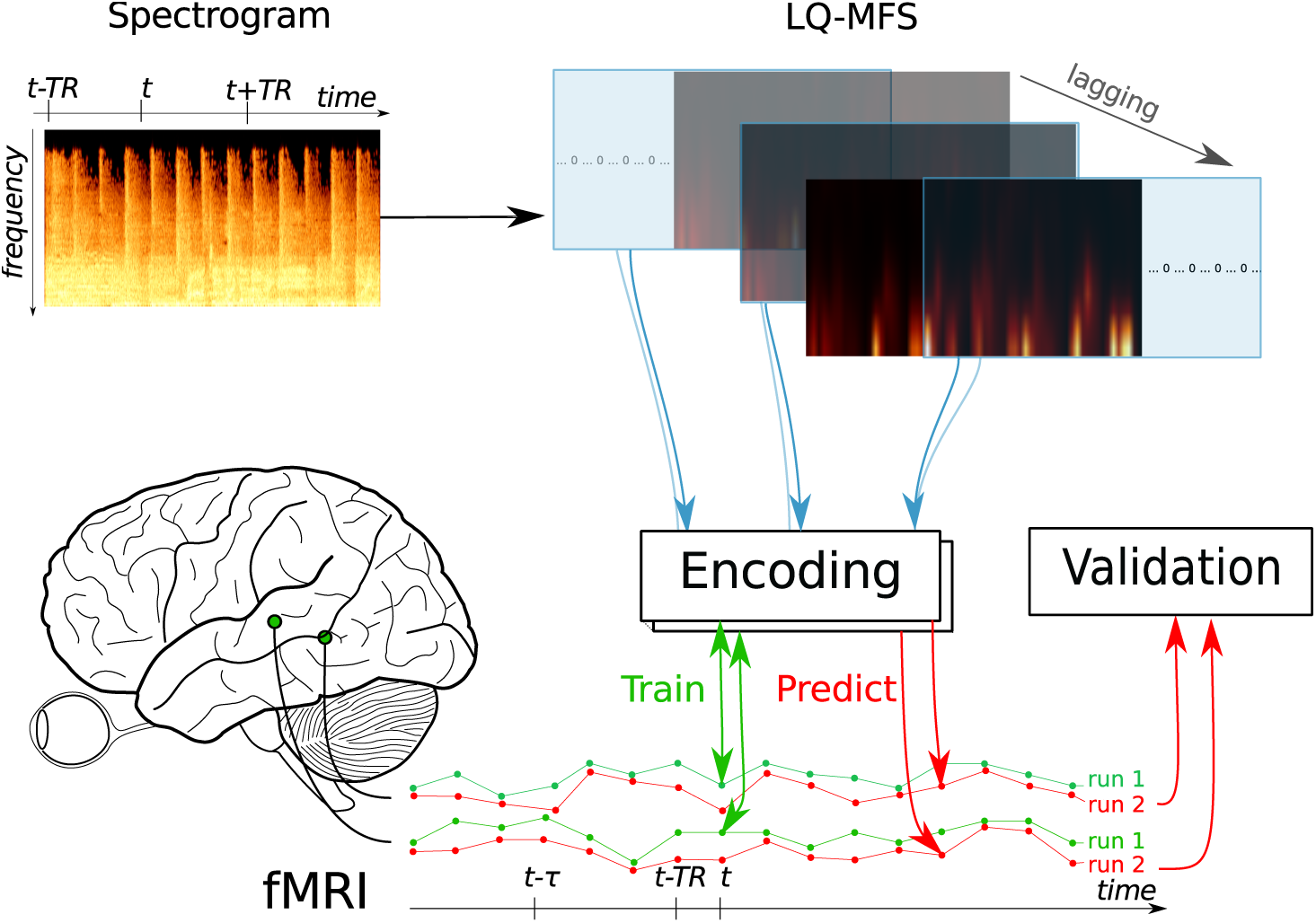
A schematic overview of the encoding process. The spectrogram for each stimulus is transformed into its low-quefrency mel-frequency spectrum (LQ-MFS). Then, the encoding features are extracted by a sliding window from the LQ-MFS. Using these features, encoding model is trained on all runs in the training set, and used to predict the BOLD activity of the left-out run. These predictions are subsequently used for validation.

The BOLD activity time-series, as well as the feature time-series, were vertically stacked, resulting in a matrix of features *X* [*N ×M*] (where *N* is the number of fMRI samples, and *M* is number of LQ-MFS coefficients, with 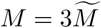) and a matrix of BOLD activity *Y* [*N ×V*] (where *V* is the number of voxels and *N* as above). This lagging of the stimulus allows us to train the encoding model to predict the fMRI time-series without explicitly modelling the BOLD response. The encoding model could then be expressed as the probability to observe the BOLD activity at time-point *t* and voxel *v*:

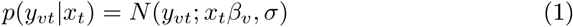

where *N* (*y*; *µ, σ*) denotes the probability density at *y* for a Gaussian with mean *µ* and standard deviation *σ*, and *β*_*v*_ is a [*M ×* 1] vector of regression coefficients specific to voxel *v*. The corresponding matrix vector equation for all voxels is

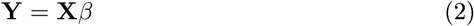

where **Y** and **X** are defined as above and *β* is a [*M × V*] matrix of regression coefficients per voxel. To reduce over-fitting, the regression-coefficients were estimated using ridge regression [Hoerl and Kennard, 1970]. Independently for each voxel, the regularization parameter *λ* with the lowest mean squared error in a generalized leave-one-out cross-validation [Golub et al., 1979] was chosen from a set of candidate values. This set was chosen so that the highest and lowest values of *λ* were only rarely selected in cross-validation. While this regularization parameter can exert a large influence on regression coefficients, and hence interpretation, we do not scrutinize its role here, since we are interested in choosing the amount of regularization that maximizes prediction accuracy. All analyses were implemented using custom code that heavily utilizes the SciPy ecosystem [Jones et al., 2014], Scikit-Learn [Pedregosa et al., 2011], Nilearn [Abraham et al., 2014], and PyMVPA [Hanke et al., 2009].

### Quality metrics

Binary retrieval accuracy and the matching score quantify encoding model performance based on the difference between predicted and observed spatial patterns that stretch across multiple voxels. They assume that each stimulus is associated with one spatial patterns, and thus do not incorporate stimulus time-courses with multiple BOLD samples. Since our stimuli encompass three BOLD samples each, we treat each BOLD sample as as an individual stimulus in both the binary retrieval accuracy and the matching score. However, we need to avoid misclassifications in the case in which predicted BOLD activity for one sample of the stimulus is closest to another sample in the same stimulus - and thus “misclassifies” the time point, but correctly classifies the stimulus. To deal with this issue, when testing each predicted BOLD sample we temporarily remove all other samples of the same stimulus from the validation set. Hence, for each BOLD sample of a stimulus we test against *N* = 25 *×* 3 *−* 2 other stimuli per run (25 stimuli with 3 BOLD samples each). Since all quality metrics are computed on the validation set in a leave-one-run-out cross-validation scheme, this leads to different number of samples used for the binary retrieval accuracy and machting score (number of samples *N* = 3 samples *×* 25 stimuli *×* 8 runs = 600), and decoding accuracy (*N* = 25 stimuli *×* 8 runs = 200 or *N* = 5 genres *×* 8 runs = 40).

### Binary retrieval accuracy

Binary retrieval accuracy [Mitchell et al., 2008] tests if an encoding model’s predictions can, on average, differentiate a stimulus’ observed BOLD activity from the BOLD activity of a decoy stimulus. Specifically, a stimulus pair is counted as succesfully classified, if the cosine similarity between predicted and observed fMRI responses is greater for the correctly matched predictions and observations than for the incorrectly matched ones (i.e. the similarity of the observed response of stimulus A with the predictions for stimulus B and vice versa). In our case, the binary retrieval measure for one predicted BOLD sample of a stimulus was computed by counting the correct matches for all (exhaustive) pair-wise combinations of all BOLD samples of all stimuli, independent of genre, while excluding other samples of the same stimulus, and then dividing by the number of combinations. This quantity was averaged across all samples of all stimuli.

### Matching score

An alternative measure of encoding performance is the correlation rank score or matching score [Santoro et al., 2014]. For each BOLD sample of a given stimulus *l*_*n*_ in the validation set, its predicted BOLD activity 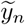 is correlated with the observed BOLD activity of every stimulus, *y*_*n*_. These correlations are then ordered, and the matching score *m*(*l*_*n*_) is

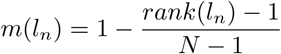

Where *rank*(*l*_*n*_) is the rank of the correlation between predicted 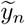 and observed *y*_*n*_ BOLD activity of *l*_*n*_, and *N* = 25 stimuli *×* 3 samples - 2 = 73 is the number of BOLD samples in the validation set excluding the two other BOLD samples from the same stimulus. Finally, the matching scores for all stimuli in the validation set are averaged.

### Decoding

Instead of testing the encoding performance, we can also test the performance of a decoder based on the individual encoding models [Naselaris et al., 2011] (Figure 2). Our goal is to obtain the probability of which music stimulus was presented from the predicted spatial activation pattern across voxels. To go from the probability of a voxel activation given stimulus features *p*(*y*_*vt*_|*x*_*t*_) to probability of stimulus features given voxel activation *p*(*x*_*t*_ |*y*_*t*_) we follow Naselaris et al. [2009] and separate our decoding scheme into two parts.

**Figure 2:**
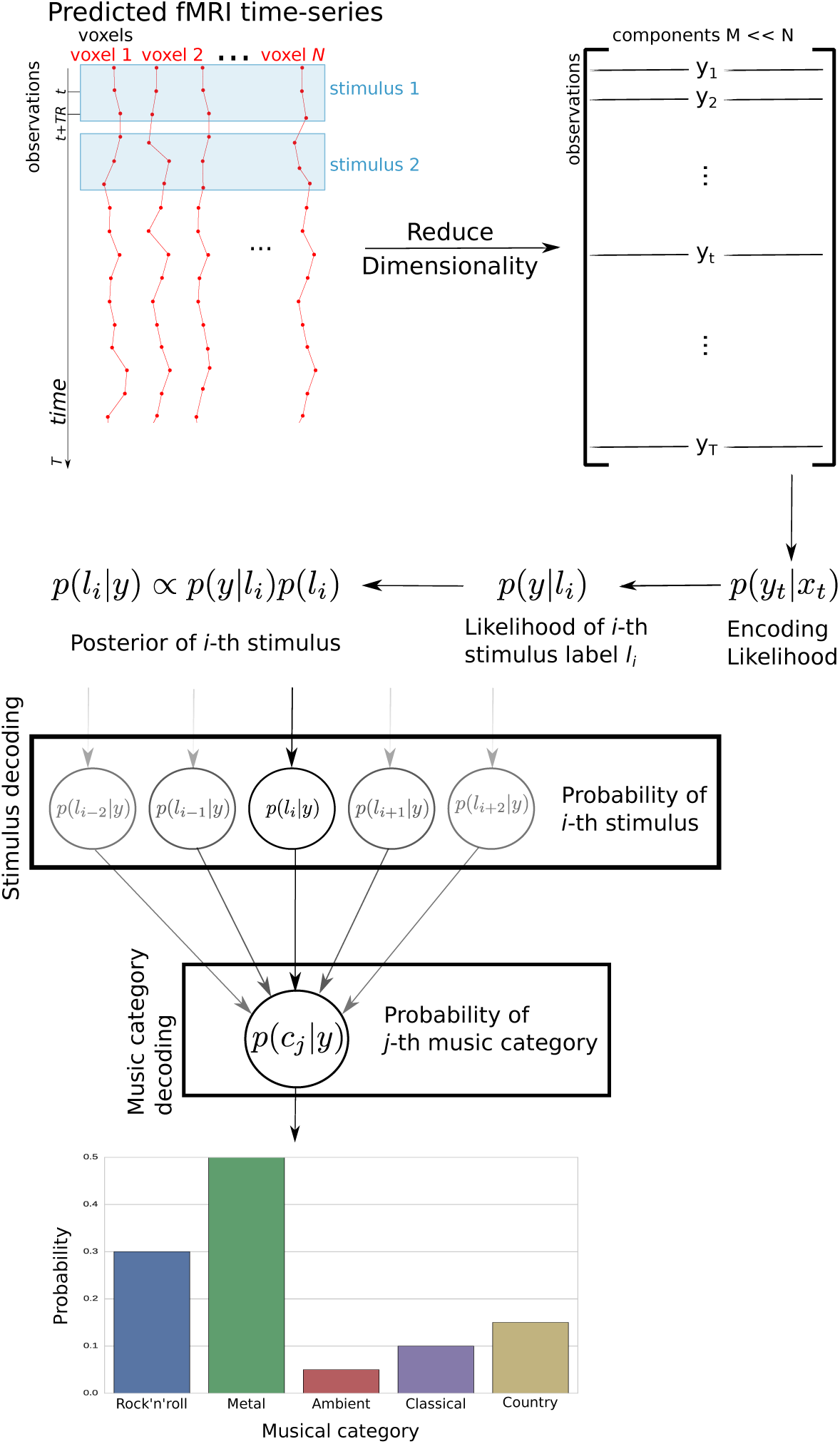
A schematic overview of the decoding of stimulus identity and stimulus music category. The predicted fMRI time series of the validation run is reduced in dimensionality by principal component analysis, and a multivariate-normal likelihood function *p*(*y*_*t*_|*x*_*t*_) is constructed. From there, the probability distribution over music stimuli and subsequently musical categories is estimated.

#### Single- to Multi-Voxel Encoding

First we condense the large number of voxel-specific encoding models into one multi-voxel encoding model. To do this we project the predicted and observed fMRI data onto the first *k* principal components of the [*N × V*] matrix of predicted BOLD activity. We choose *k* via cross-validation to estimate the number of principal components that maximize the decoding accuracy on the training set for each participant. As in Naselaris et al. [2009] we construct the *k*-dimensional multivariate normal probability density function *p*(*y*_*t*_|*x*_*t*_) to obtain a likelihood function across voxels.

#### Multi-Voxel Encoding to Decoding

We now express this likelihood in terms of the label of the music stimulus, instead of its LQ-MFS features. We use the simplifying assumption that the BOLD activity is influenced by the music stimuli only through their LQ-MFS coefficients *x*, and - given that each music stimulus was associated with only three (lagged) LQ-MFS representations - the likelihood to observe a given triple of consecutive *y* for a specific music stimulus *l*_*n*_ is *p*(*y*|*l*_*n*_) ∝ *p*(*y*_*t−*1_|*x*_1_)*p*(*y*_*t*_|*x*_2_)*p*(*y*_*t*+1_|*x*_3_) where *t* is the sample 6 seconds after the start of the music stimulus, *x* are the three LQ-MFS feature vectors of this stimulus, and *n* = 1..25 denotes the stimulus. For a given triple of consecutive BOLD activity *y* from the same stimulus, we can now estimate the probability distribution over music stimuli *p*(*l*_*n*_|*y*) by using Bayes’ rule: *p*(*l*_*n*_|*y*) ∝ *p*(*y*|*l*_*n*_)*p*(*l*_*n*_). Since each stimulus was presented exactly once, it has an uniform prior distribution with 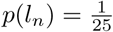. The mode of this distribution is the most probable presented stimulus given the data. Additionally we decode the musical genre of the presented stimulus given the observed BOLD activity as the mode of 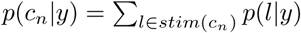 where *stim*(*c*_*n*_) are the labels of the stimuli belonging to the genre *c*_*n*_.

### Voxel selection

We varied the number of voxels used in the analysis, both for 3T and 7T fMRI data, and selected which voxels to keep by two different criteria. Both criteria were based on voxel characteristics in the training set only.

#### Selection by stability

Mitchell et al. [2008] selected the 500 most stable voxels for their analysis. For a single voxel, each run can be represented as a vector of BOLD activity, where each entry is associated with one stimulus.

A voxel’s “stability score” is then the average (pair-wise) correlation between the vectors of the eight runs for all combinations of runs. Hence, this criterion selects voxels with consistent activation for each stimulus across runs.

#### Selection by *r*^2^

As we are interested in encoding performance, we can use the quality of predictions of each voxel’s encoding model as a selection criterion. We compute the coefficient of determination *r*^2^ for each voxel-specific encoding model in the training set. Using this criterion selects voxels whose activity can be explained best by an LQ-MFS-based encoding model.

## Results

We trained voxel-wise encoding models to predict voxel activity using a LQ-MFS representation of music stimuli in 3T and 7T fMRI data. First, we evaluate each voxel’s encoding model individually. Figure 3 shows *r*^2^ values from an eight fold leave-one-run-out cross-validation of the 10000 voxels with highest *r*^2^ averaged across participants. While voxel-wise encoding models show higher peak *r*^2^ in 3T than in 7T in the best performing models (left side of the figure), voxel-wise encoding models show higher *r*^2^ in 7T than in 3T for all other voxels.

**Figure 3:**
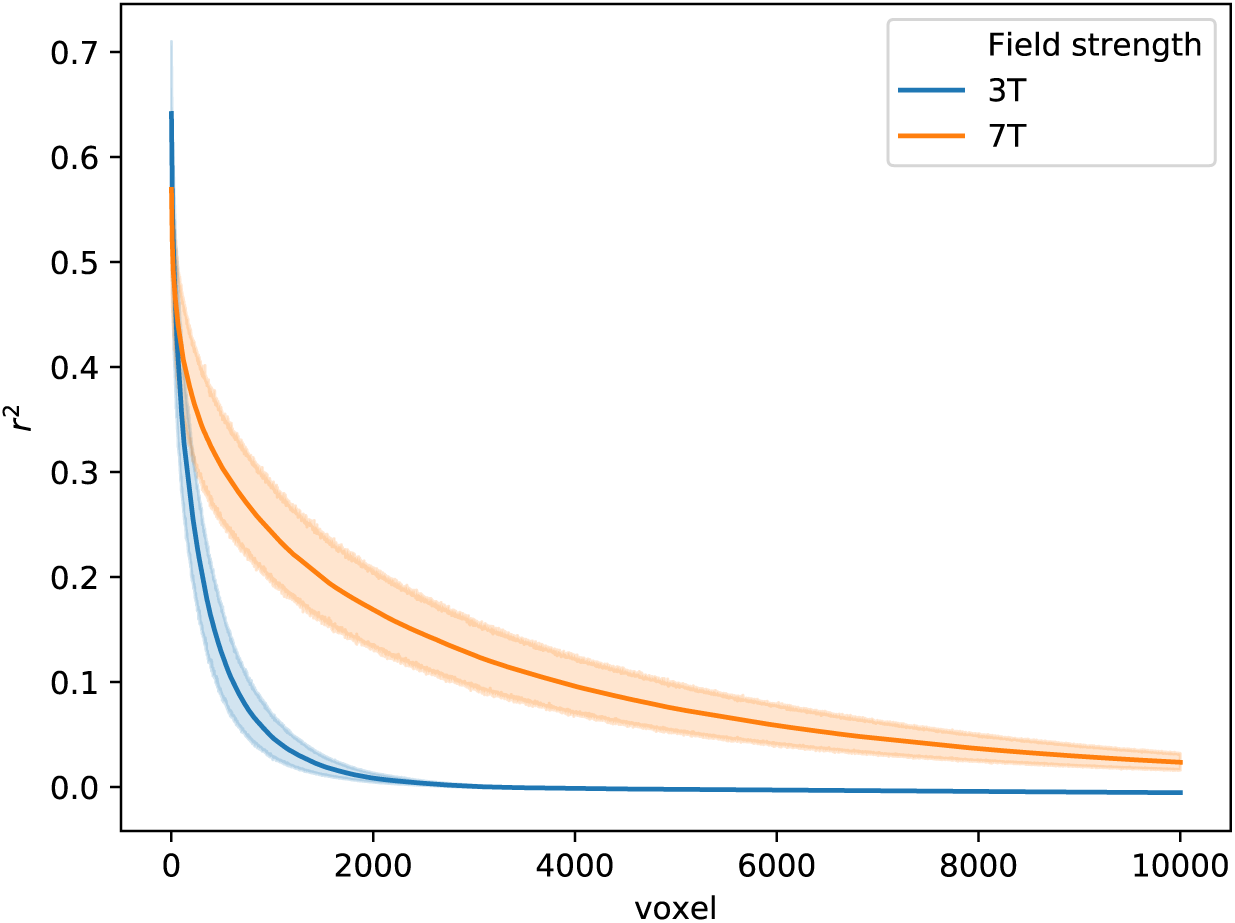
Voxel-wise *r*^2^ between predicted and observed BOLD activity for each left-out run, averaged across participants and runs with 95% confidence interval. Shown here are the 10000 voxels with highest *r*^2^ for 3T and 7T.

Next, we evaluate sets of voxel-wise encoding models for 3T and 7T using several quality metrics. We varied the number of voxels used in the analysis and how they were selected, both for 3T and 7T fMRI data, and compared the resulting differences in three quality metrics of encoding models. To differentiate between effects of voxel number and overall volume of the voxels, which differs in 3T and 7T, we show the results as a function of the number of voxels, as well as overall volume of these voxels. To illustrate the extent of 10000 selected voxels for different field strengths and selection criteria, we show for each voxel the proportion of validation sets across all participants in which it was selected in Figure S1. In both 3T and 7T the voxels that are most often selected are situated in primary and secondary auditory areas.

### Binary retrieval accuracy

Figure 4 shows differences in binary retrieval accuracy for different numbers of voxels, selection strategies, and field strength. For both 3T and 7T data, and both selection strategies, the maximum binary retrieval accuracy is achieved with relatively low number of voxels (3T and *r*^2^: 250 voxels, 6.75 *cm*^3^ volume, 7T and *r*^2^: 500 voxels, 0.68 *cm*^3^ volume, 3T and stability selection: 500 voxels, 13.5 *cm*^3^ volume, 7T and stability selection: 1000 voxels, 2.7 *cm*^3^ volume) and decreases the more voxels are included. For both stability selection criteria, 3T outperforms or is equal to 7T across all numbers of voxels. With regard to the voxel selection strategies, selection by *r*^2^ outperforms or is equal to selection by stability across all numbers of voxels (Figure S2).

**Figure 4:**
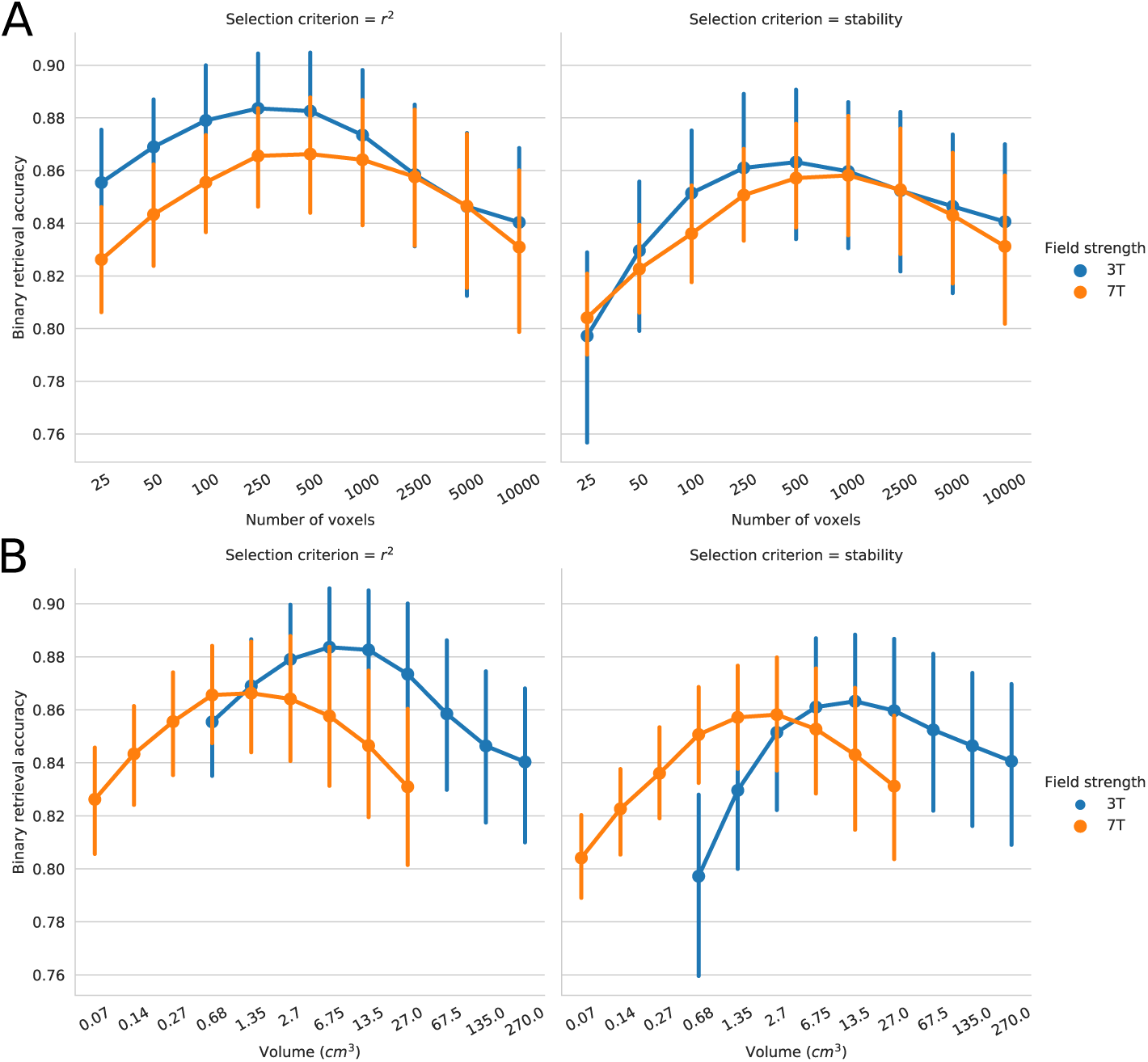
**A** Mean binary retrieval accuracy as a function of the included number of voxels for 3T and 7T, for stability- and *r*^2^-based voxel selection. Error bars denote the bootstrapped 95% confidence interval of the mean. The mean is taken over binary retrieval accuracies of eight runs for each of the 19 participants. **B** Mean binary retrieval accuracy as a function of the overall volume of the included voxels for 3T and 7T, for stability- and *r*^2^-based voxel selection.

### Matching score

Figure 5 shows differences in matching score for different numbers of voxels, selection strategies, and field strength. Matching scores in both 3T and 7T data peak at high numbers of voxels (3T and *r*^2^: 1000 voxels, 27 *cm*^3^ volume, 7T and *r*^2^: 5000 voxels, 13.5 *cm*^3^ volume, 3T and stability selection: 500 voxels, 13.5 *cm*^3^ volume, 7T and stability selection: 5000 voxels, 13.5 *cm*^3^ volume). Indexing by the overall volumes these voxels encompass reveals that the matching score peaks for 3T and 7T at (for stability selection) or close to (for selection by *r*^2^) the same volume. For both stability selection and selection by *r*^2^, 3T outperforms 7T for the same number of voxels, except for the largest two numbers of voxels, and - in terms of the volume the voxels take - 7T outperforms 3T for volumes up to 2.7 *cm*^3^ and 3T outperforms 7T for volumes beyond 27 *cm*^3^. Furthermore, for 3T data, selection by stability and selection by *r*^2^ perform similarly, while selection by *r*^2^ outperforms selection by stability consistently in 7T data (Figure S3).

**Figure 5:**
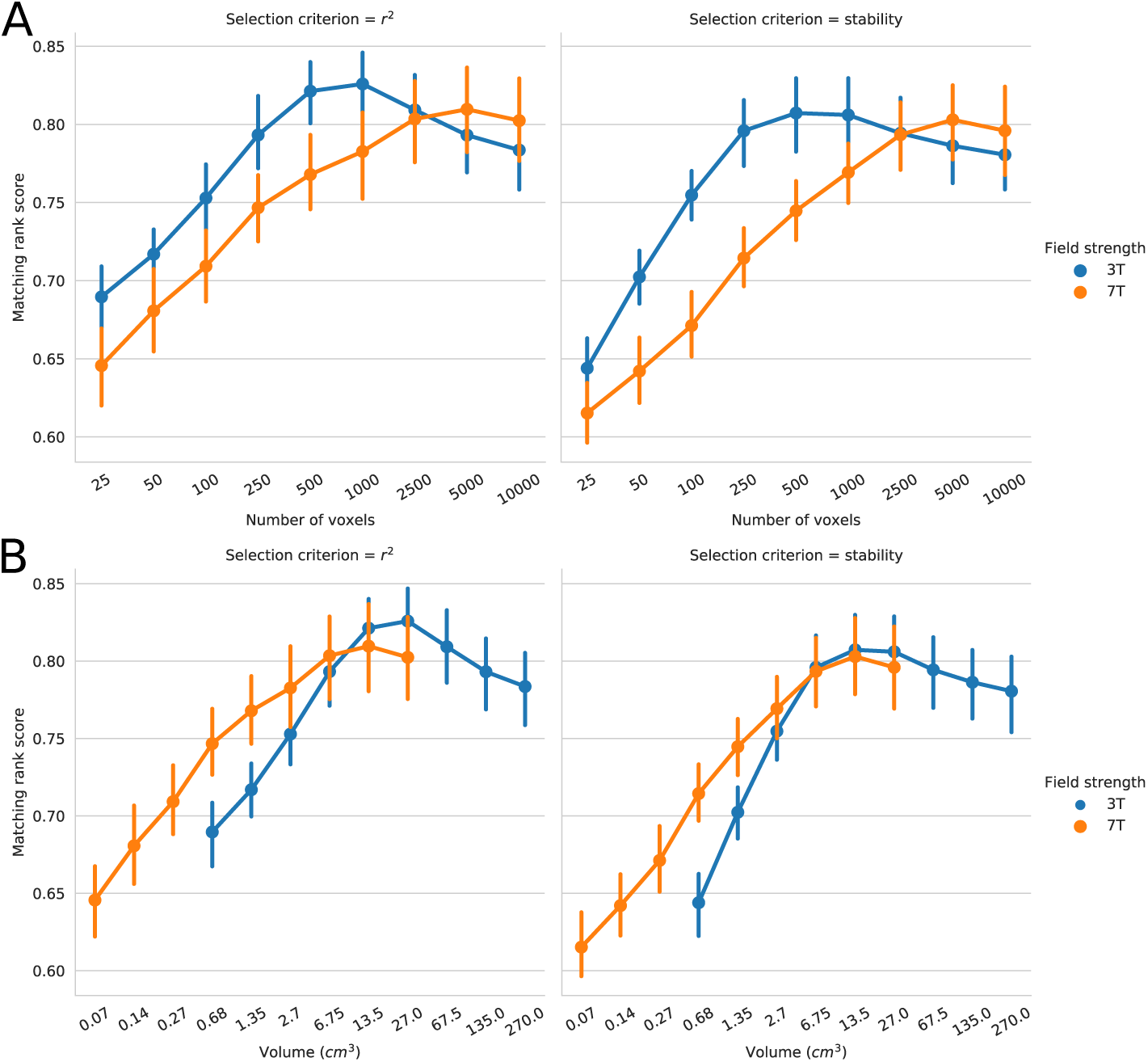
**A** Mean matching rank score as a function of the included number of voxels for 3T and 7T, for stability- and *r*^2^-based voxel selection. Error bars denote the bootstrapped 95% confidence interval of the mean. The mean is taken over binary retrieval accuracies of eight runs for each of the 19 participants. **B** Mean matching rank score as a function of the overall volume of the included voxels for 3T and 7T, for stability- and *r*^2^-based voxel selection.

### Decoding accuracy

Figure 6 shows the accuracy of decoding each individual stimulus (chance level 0.04) for different numbers of voxels, selection strategies, and field strength. 3T consistently outperforms 7T in decoding individual stimuli: if voxels are selected by stability, 3T outperforms 7T for all numbers of voxels, if voxels are selected by *r*^2^, 3T outperforms 7T for all but the two highest numbers of voxels. For 7T data, the number of voxels matters little, however, for 3T data, optimal decoding accuracy is reached at a relatively low number of voxels (250 voxels for selection by *r*^2^ and selection by stability, 6.75 *cm*^3^ volume), and subsequently decreases if more voxels are included.

**Figure 6:**
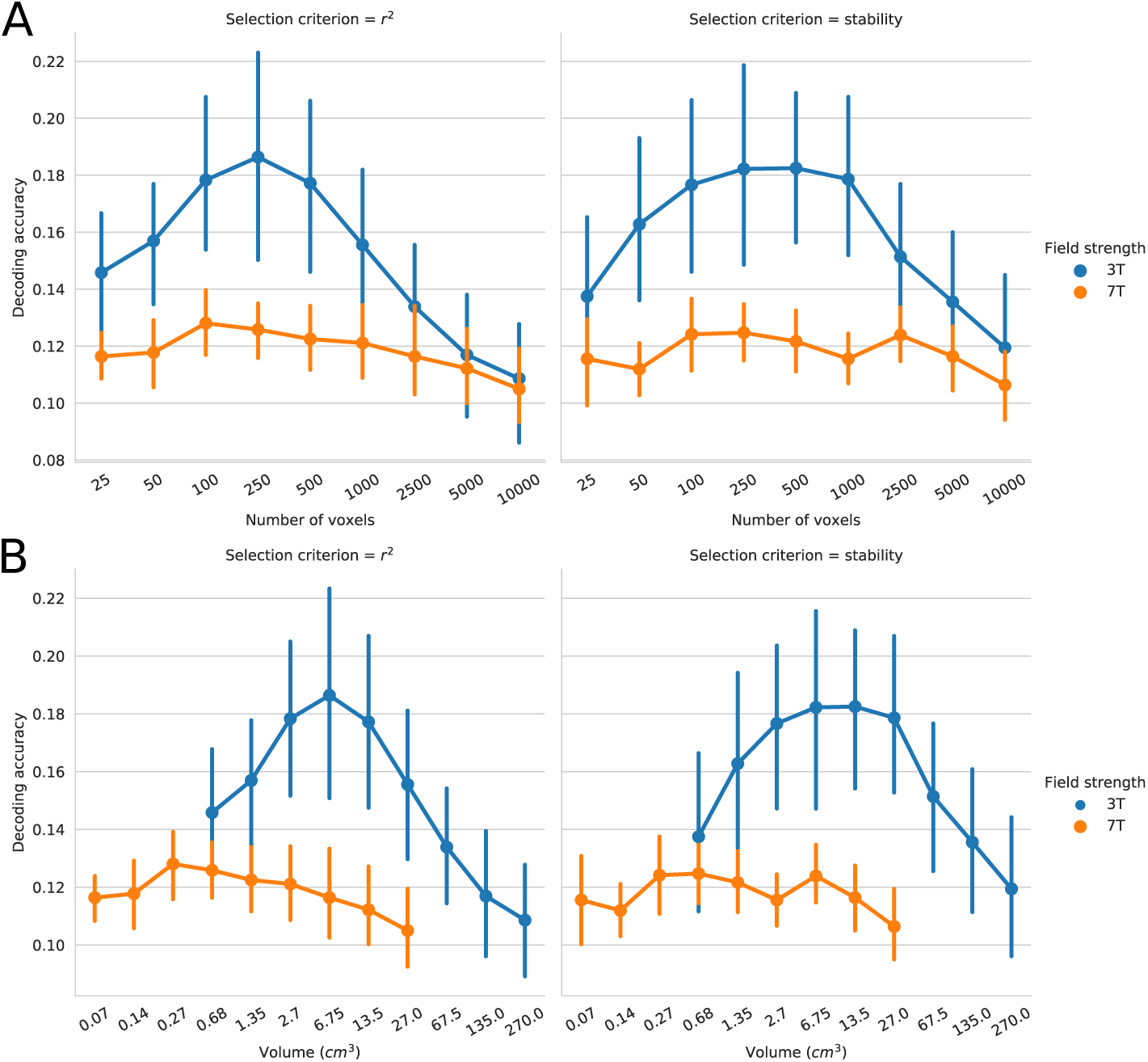
**A** Mean decoding accuracy of individual music stimuli as a function of the included number of voxels for 3T and 7T, for stability- and *r*^2^-based voxel selection. Error bars denote the bootstrapped 95% confidence interval of the mean. The mean is taken over decoding accuracies of eight runs for each of the 19 participants. Chance level is 0.04. **B** Mean decoding accuracy of individual music stimuli as a function of the overall volume of the included voxels for 3T and 7T, for stability- and *r*^2^-based voxel selection. Chance level is 0.04.

Instead of decoding individual stimuli, we can also decode the musical genre of each individual stimulus. Figure 7 shows the accuracy of decoding music genre (chance level 0.20) for different numbers of voxels, selection strategies, and field strength. Here a different pattern emerges: decoding accuracy is higher for 7T data than for 3T data, across all numbers of voxels and selection criteria. Potentially, this suggests that 7T BOLD activity contains different information than 3T. For both selection strategies, 7T shows higher maximum genre decoding accuracy for high numbers of voxels, while 3T shows higher maximum genre decoding accuracy for lower numbers of voxels.

**Figure 7:**
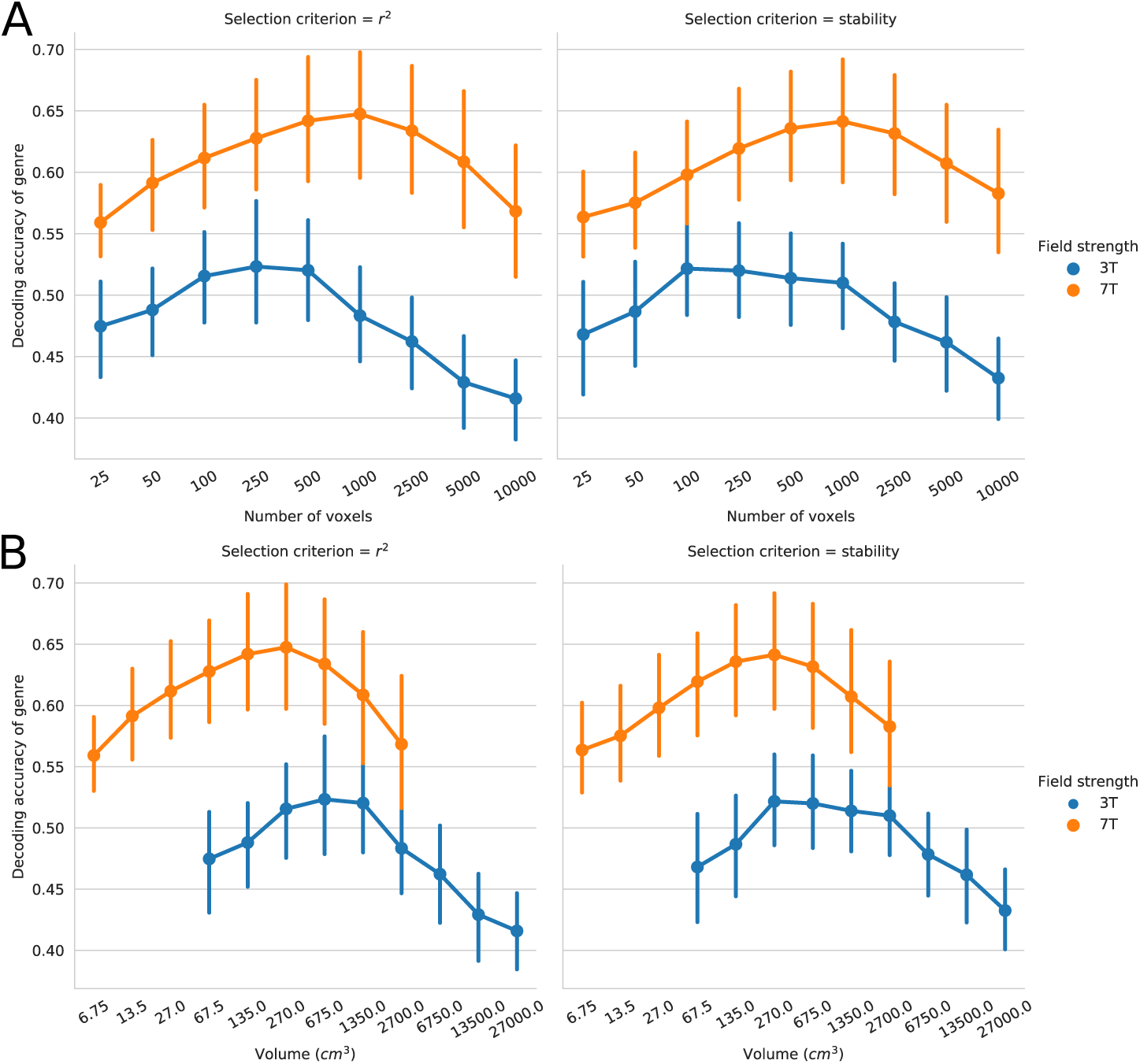
**A** Mean decoding accuracy of music category as a function of the included number of voxels for 3T and 7T, for stability- and *r*^2^-based voxel selection. Error bars denote the bootstrapped 95% confidence interval of the mean. The mean is taken over decoding accuracies of eight runs for each of the 19 participants. Chance level is 0.2. **B** Mean decoding accuracy of music category as a function of the overall volume of the included voxels for 3T and 7T, for stability- and *r*^2^-based voxel selection. Chance level is 0.2.

Next, we want to answer the question which musical genres can be best predicted, and if those differ between 3T and 7T data. For this we compute the confusion matrix from the out-of-sample predictions of all classifiers: Rows denote the stimulus’ true genre, columns denote the predicted genre. Each cell contains the count - or proportion - of occurences of this combination of true and predicted genres. Maximum classification accuracy leads to a diagonal confusion matrix.

Figure 8 shows the confusion matrices averaged across numbers of voxels for different field strength and selection criteria. The occurences are normalized per row, and their mean proportion across numbers of voxels are indicated in each cell. All confusion matrices show a higher misclassification between genres that contain speech (country, rock’n’roll and heavy metal) and between instrumental genres (ambient and symphonic), while very few genres that contain speech are incorrectly classified as instrumental or vice versa. The increased genre decoding accuracy in 7T is thus due to a better discrimination between the two instrumental genres ambient and symphonic - and between the three vocal genres - country, rock’n’roll, and heavy metal.

**Figure 8:**
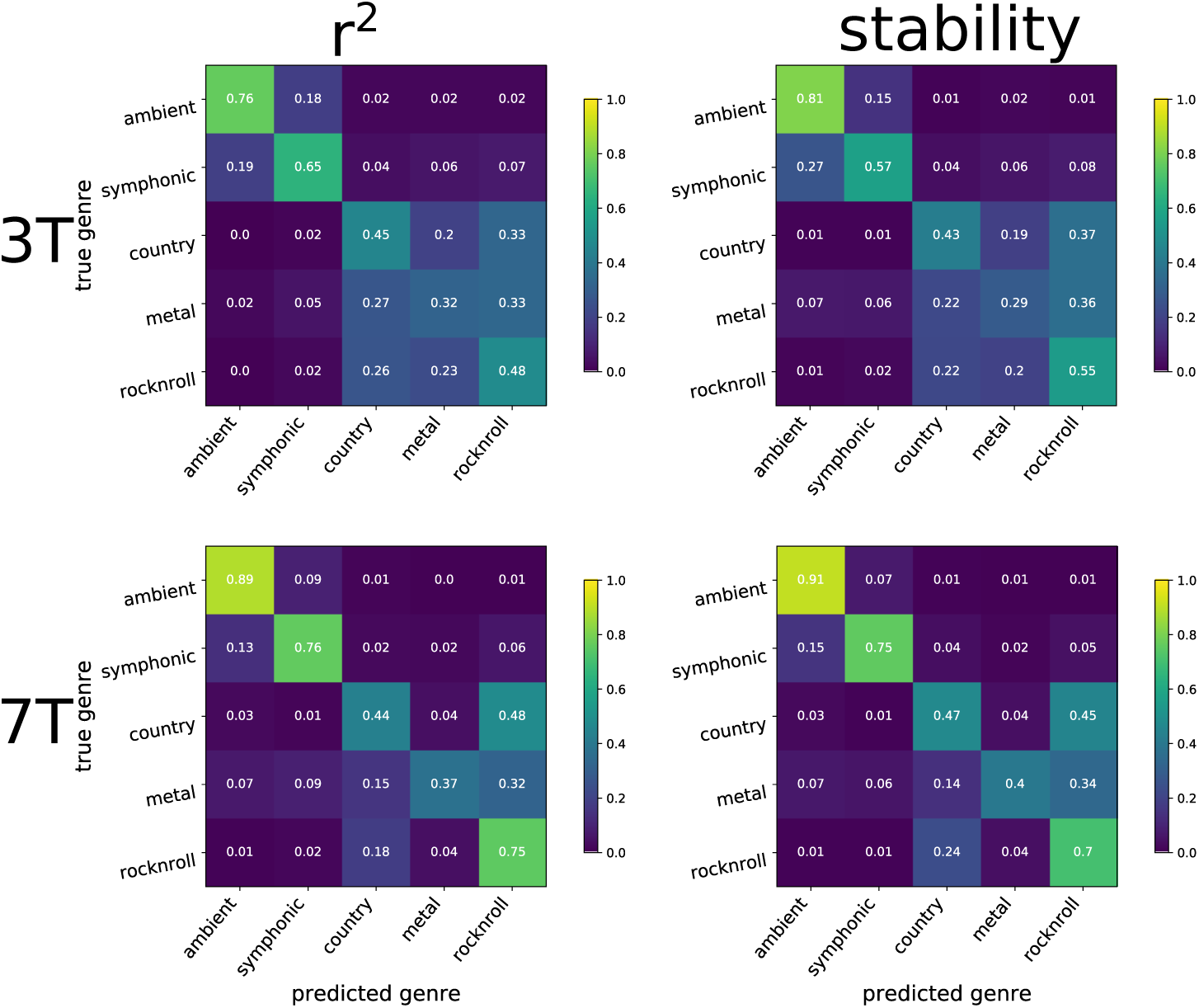
Confusion matrices for genre decoding, separated by field-strength and voxel selection. Each confusion matrix is averaged across the different number of voxels, participants and runs.

## Discussion

Our results show that all three quality metrics vary with the number of voxels and voxel selection strategies in both low and high field strengths. We demonstrate that the patterns of this variation differ between quality metrics and that field strength, voxel selection, and voxel number all affect these metrics differently.

We believe these results emphasize the significance of an often under-appreciated aspect of computational models in neuroimaging: near arbitrary choices in the data analysis - like the number of voxels used or how they are selected - have a considerable impact on the performance evaluation of voxel-wise encoding models. Furthermore, the impact of these choices is inconsistent across the considered validation strategies. The effects of these parameters are rarely known a-priori, and each choice of a specific value can often be easily justified, yet this undisclosed flexibility leads to considerable researcher degrees-of-freedom [Simmons et al., 2011, Hong et al., 2019] that will bias the comparison between different voxel-wise encoding models and hinder generalization of findings across individual studies.

In light of the substantial effects individual parameters exert on validation strategies, it does not surprise that one of our results - a higher matching score in 7T than in 3T BOLD fMRI data - constrasts a previous study [Santoro et al., 2014], which found higher matching scores for encoding natural sounds in 7T than in 3T, although using a different number of stimuli in the two conditions.

However, in spite of these variabilities, we can abstract several recommendations for applying and validating voxel-wise encoding models.

First, it is likely that selecting voxels by the quality of its encoding model, such as the *r*^2^ score, will outperform a selection of voxels based on their stable response to the stimulus - however, this difference diminishes for large numbers of voxels. One reason that this effect is especially prominent when only few voxels are included could be the quickly diminishing returns for higher number of voxels: voxels in which an encoding model performs well are already included, and each additional voxel decreases the joint encoding performance. Interestingly, this pattern differs in the matching score, where performance in 7T is worse for the smallest number of voxels and increases when more voxels are included. In fact, this holds true for selection by both *r*^2^ and a stability criterion, and thus differentiates the matching score from both binary retrieval accuracy, as well as decoding accuracy. This contrasts especially with binary retrieval accuracy, results from a previous study indicate that low numbers of voxels lead to high binary retrieval accuracy [Hoefle et al., 2018]. Our results seem to corroborate this finding, suggesting that binary retrieval accuracy, in contrast to the matching score, is sensitive to the inclusion of worse performing voxels.

Second, the number of voxels used has a large effect on all quality metrics, and to compare two sets of voxel-wise encoding models even on the same stimuli, one has to control for them. For most quality metrics, this control is most easily exercised by fixing the number of voxels directly, however, in case of the matching score, it can be more meaningful to fix the overall volume the two sets of encoding models encompass.

Third, while higher field strength might lead to better individual encoding models, it might not improve their decoding accuracy. We show that voxel-wise encoding models in 3T fMRI data outperform voxel-wise encoding models in 7T fMRI data in all but one quality metric. This indicates that high field strength does not provide more information per se, since stimuli can’t be differentiated better in general, in spite of the higher *r*^2^ in most voxels in 7T. However, our key insight is, that higher field strength can decrease the decoding accuracy for some labels of the stimulus, i.e. the decoding of individual stimuli, yet increase the decoding accuracy for other labels, i.e the decoding of the musical genre of individual stimuli. Thus, one has to exercise caution, when choosing which label to decode from a set of encoding models. One reason for this, could be that training encoding models on stimulus features can make stimulus labels somewhat arbitrary. Two similar stimuli can lead to similar and accurate predictions of BOLD activity, and hence a higher *r*^2^, yet this similarity could decrease a quality measure that is based on the correct discrimination between individual stimuli.

While we believe that these three recommendations can guide neuroimaging researchers, to conclusively assess if these recommendations generalize, more data are needed, since the results in this study are based on only two fMRI datasets encompassing the same auditory stimuli recorded in 3- and 7T. This facilitates a comparison between different field strengths, yet, further work is needed to show that these conclusions hold beyond this set of stimuli, our chosen stimulus representation, and even beyond the auditory domain.

## Author contributions

MB performed the analysis and wrote the manuscript. JSG contributed to the manuscript. JWR contributed to the manuscript. MH contributed to the manuscript.

## Acknowledgements

We are grateful to Michael Casey and the musicians. We are also thankful for Cristiano Micheli’s helpful feedback on an earlier version of this manuscript.

## Supplementary information

**Figure S1:**
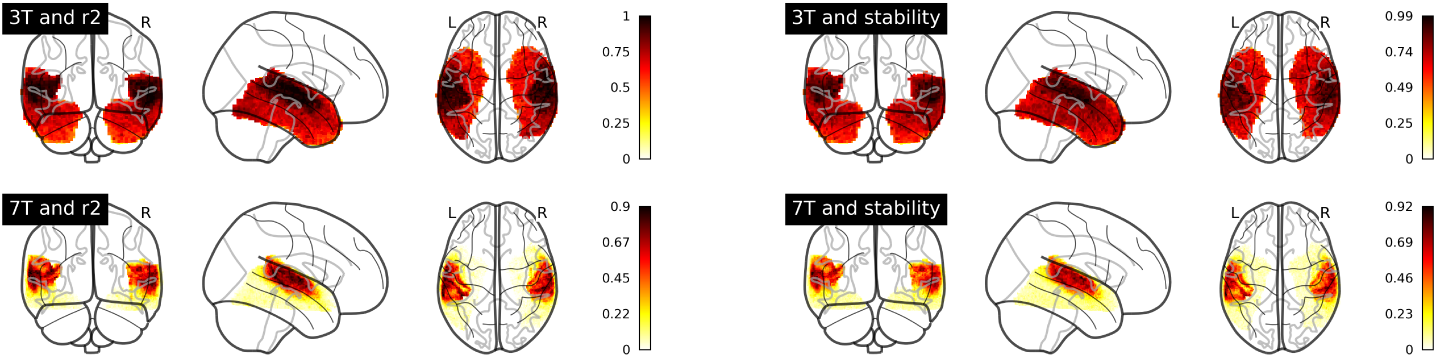
Glassbrains showing the proportion of how often a voxel was included across all participants and all validation sets for different field strengths and voxel selection criteria.

**Figure S2:**
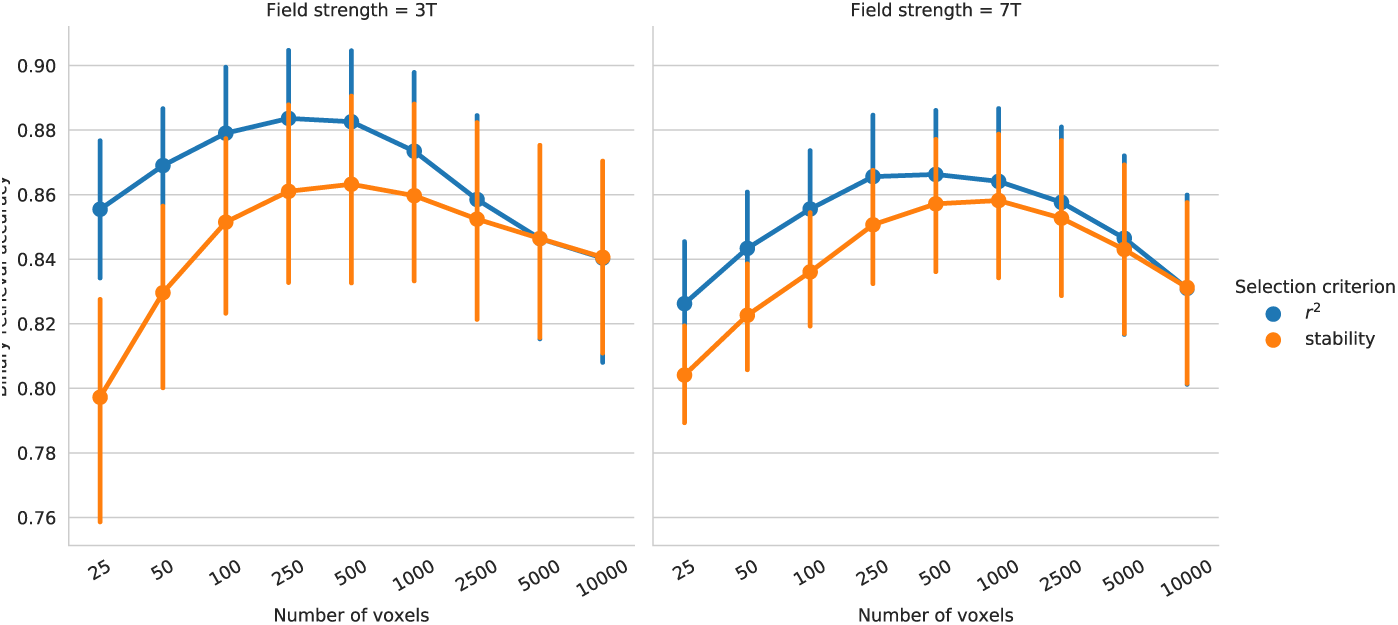
**A** Mean binary retrieval accuracy as a function of the included number of voxels for 3T and 7T, for stability- and *r*^2^-based voxel selection. Error bars denote the bootstrapped 95% confidence interval of the mean. The mean is taken over binary retrieval accuracies of eight runs for each of the 19 participants.

**Figure S3:**
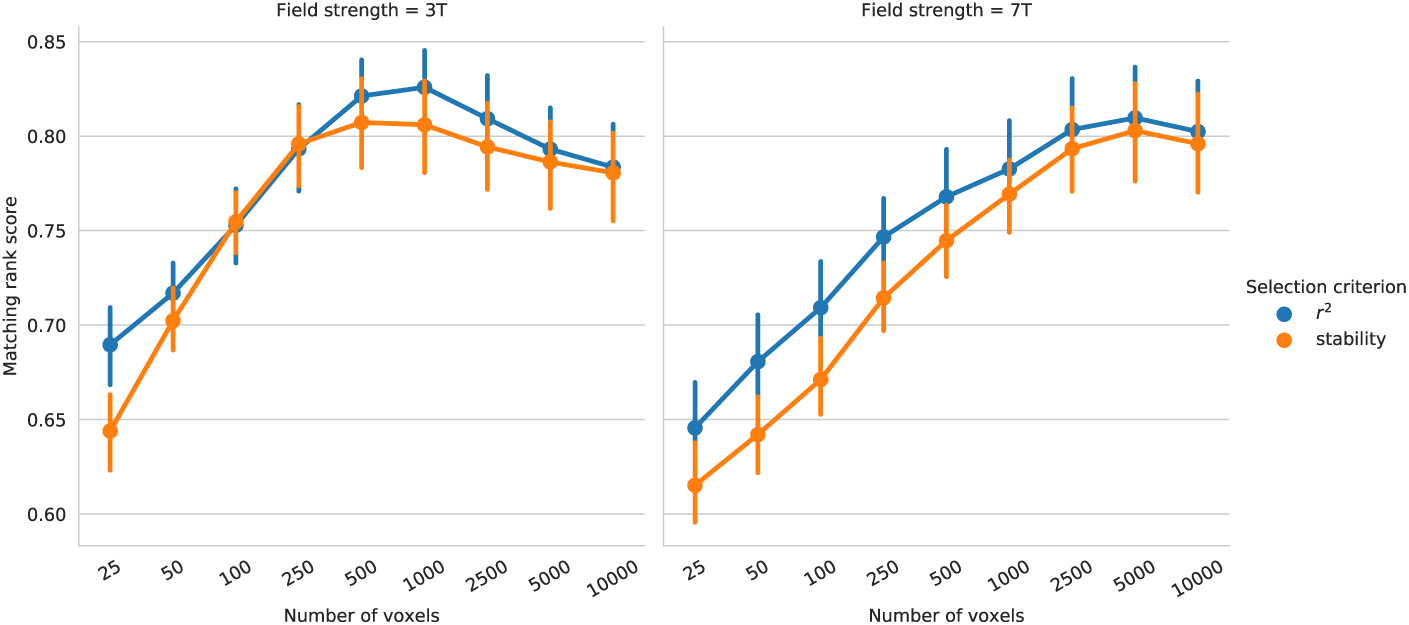
**A** Mean matching rank score as a function of the included number of voxels for 3T and 7T, for stability- and *r*^2^-based voxel selection. Error bars denote the bootstrapped 95% confidence interval of the mean. The mean is taken over binary retrieval accuracies of eight runs for each of the 19 participants.

**Figure S4:**
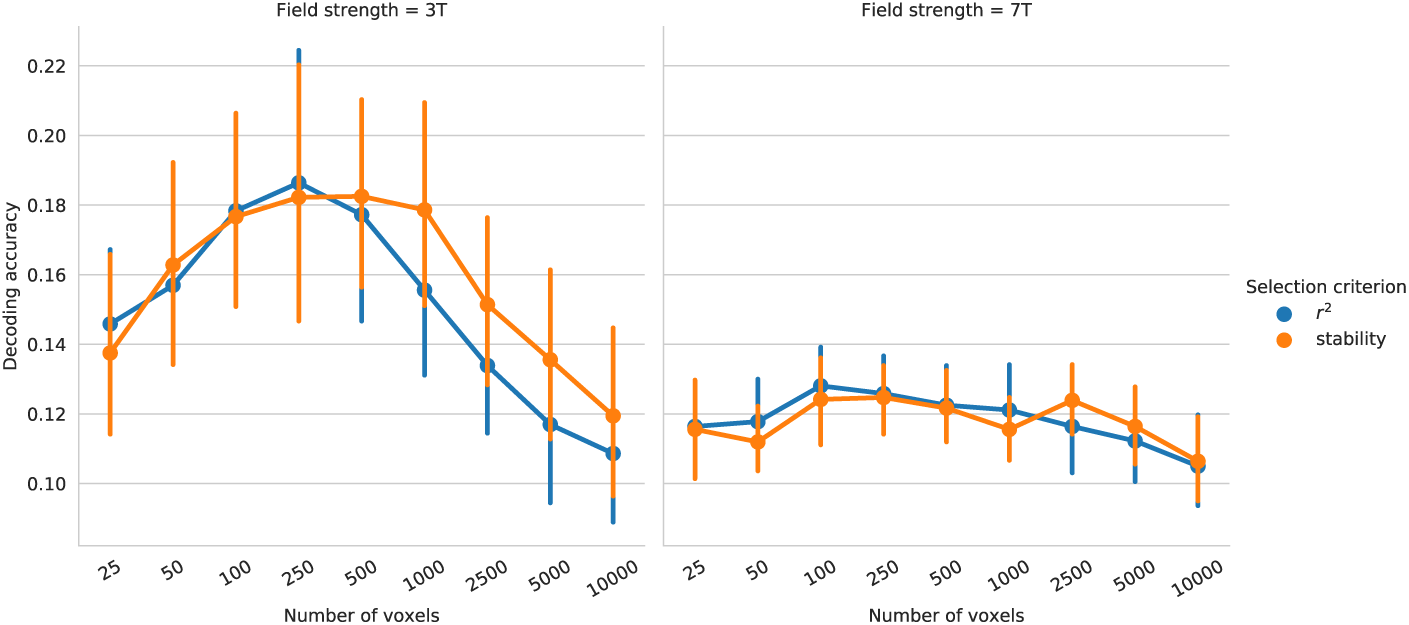
**A** Mean decoding accuracy of individual music stimuli as a function of the included number of voxels for 3T and 7T, for stability- and *r*^2^-based voxel selection. Error bars denote the bootstrapped 95% confidence interval of the mean. The mean is taken over decoding accuracies of eight runs for each of the 19 participants. Chance level is 0.04.

**Figure S5:**
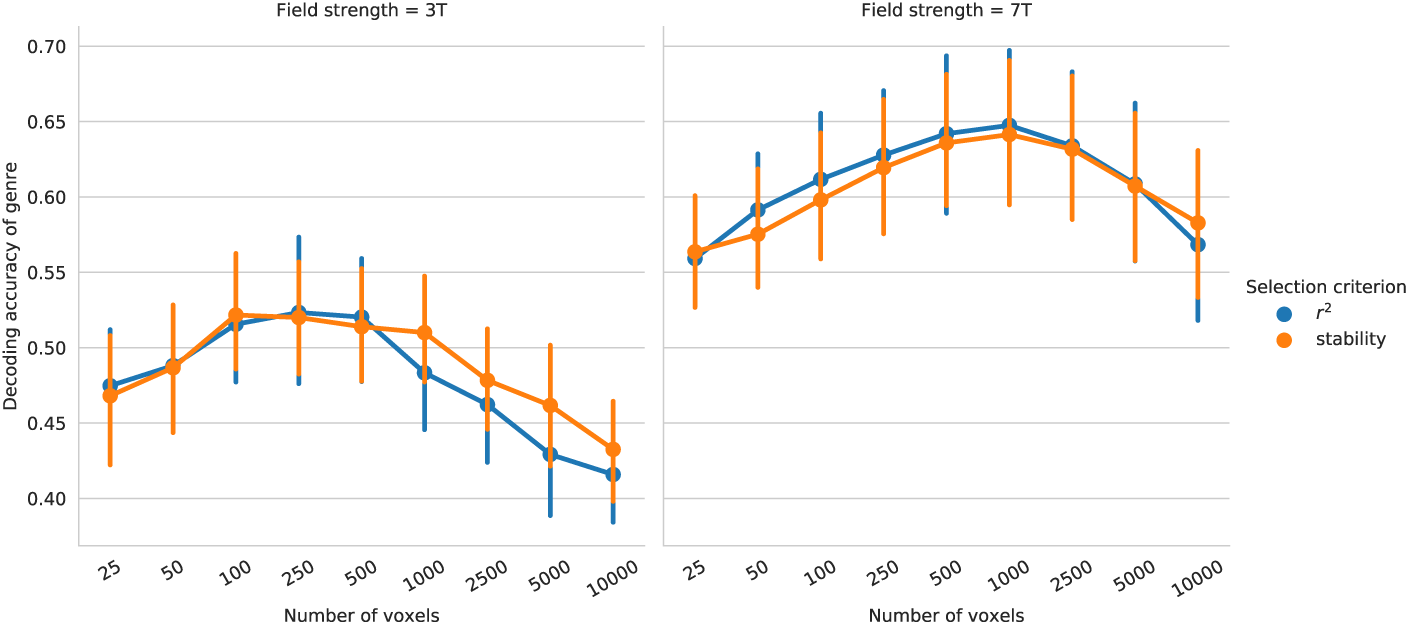
Mean decoding accuracy of music category as a function of the included number of voxels for 3T and 7T, for stability- and *r*^2^-based voxel selection. Error bars denote the bootstrapped 95% confidence interval of the mean. The mean is taken over decoding accuracies of eight runs for each of the 19 participants. Chance level is 0.2.

